# Optimality of extracellular enzyme production and activity in dynamic flux balance modeling

**DOI:** 10.1101/2021.11.01.466736

**Authors:** Michael Quintin, Ilija Dukovski, Jennifer Bhatnagar, Daniel Segrè

## Abstract

In microbial communities, many vital metabolic functions, including the degradation of cellulose, proteins and other complex macromolecules, are carried out by costly, extracellularly secreted enzymes. While significant effort has been dedicated to analyzing genome-scale metabolic networks for individual microbes and communities, little is known about the interplay between global allocation of metabolic resources in the cell and extracellular enzyme secretion and activity. Here we introduce a method for modeling the secretion and catalytic functions of extracellular enzymes using dynamic flux balance analysis. This new addition, implemented within COMETS (Computation Of Microbial Ecosystems in Time and Space), simulates the costly production and secretion of enzymes and their diffusion and activity throughout the environment, independent of the producing organism. After tuning our model based on data for a *Saccharomyces cerevisiae* strain engineered to produce exogenous cellulases, we explored the dynamics of the system at different cellulose concentrations and enzyme production rates. We found that there are distinct rates of constitutive enzyme secretion which maximize either growth rate or biomass yield. These optimal rates are strongly dependent on enzyme kinetic properties and environmental conditions, including the amount of cellulose substrate available. Our framework will facilitate the development of more realistic simulations of microbial community dynamics within environments rich in complex macromolecules, with applications in the study of soil and plant-associated ecosystems, and other natural and engineered microbiomes.

**Importance:** Many organisms - including soil, marine and human-associated bacteria and fungi - perform part of their metabolic functions outside of the boundary of the cell, through the secretion of extracellular enzymes that can diffuse and facilitate reactions independently of the organism that produced them. In order to better understand and predict microbial ecosystems, it would be helpful to create mathematical models incorporating these extracellular reactions within simulations of metabolism at the whole-cell level. In this paper we demonstrate the implementation of such a methodology and apply it to study a cellulase-secreting yeast. This work will be useful for a number of microbial ecology applications, including modeling of microbiome dynamics, engineering of bioproducts (e.g. biofuels) from plant biomass through synthetic communities or modified organisms, and testing of basic ecological hypotheses about the balance between cost and benefits of the production of common goods in microbial communities.

## Introduction

The secretion and activity of extracellular enzymes is a fundamental component of the metabolic processes of microbes and microbial communities [1–5]. These enzymes can perform a variety of functions, from the degradation of complex carbohydrate polymers, such as glycan and cellulose, to the hydrolysis of proteins and oxidation of aromatic molecules. Both the enzymes themselves and the products of the extracellular reactions they catalyze can change dynamically in the environment, modulated by diffusion, stability, and kinetic properties [6,7]. Moreover, both enzymes and molecular products constitute common goods available for all species in the community [8], posing unresolved physiological and ecological questions on the possible strategies that individual organisms may have evolved to manage these processes and interact with each other [3,9].

While the molecular, cellular and ecological properties of extracellular enzyme systems can vary greatly across different organisms and specific functions, some common fundamental questions arise in different contexts. One question is how each organism controls the expression and secretion of extracellular enzymes towards efficient growth [5,10]. A number of enzymes are produced in response to the presence of their substrate (either simple or complex [7]) or reaction products [11]. In some cases these enzymes are suppressed by an excessive abundance of the catabolite they serve to make available [12] or other preferred carbon sources [13][14]. Other extracellular enzymes are constitutively expressed, allowing a baseline rate of catalysis, with further expression possibly stimulated by the catalytic product [15]. Extracellular enzyme activities can also be sensitive to environmental conditions such as temperature, CO_2_ concentrations, and precipitation [16], as well as to quorum sensing [17]. A second question is how the molecules made available through the breakdown of biopolymers by extracellular enzymes affects different organisms in a community, including enzyme producers who carry the energetic burden of synthesizing proteins that benefit other species as well. This aspect has been extensively studied, for example in soil and litter-degrading fungal communities [18,19], where extracellular enzymes play an important role in temporal succession and ecosystem-level metabolic functions of decomposer microorganisms [6,7,18,20,21]. Similar processes are also important in the degradation of specific nutrients in the gut of humans and animals [1,22,23]. A third question is whether a quantitative mechanistic understanding of the costs and benefits of different microbial strategies for extracellular enzyme secretion can help in the design of synthetic communities, e.g. towards the construction of engineered consortia capable of pharmaceutical or biofuel production [24–26], or bioremediation of industrial waste [27].

The costs and benefits of intracellular enzyme production have been studied through experimentally testable mathematical models aimed at quantifying a hypothetical optimal expression level [28–30]. Similarly, according to economic principles [31], one would expect that an extracellular enzyme should only be produced when it can provide nutrients and energy whose benefit to the organism exceed the enzyme production cost, especially given that the secreted proteins cannot be internally recycled by the organism [5,7]. A number of papers have addressed these questions using mathematical models, generally based on systems of differential equations that describe key properties of the organism and its metabolic activity [3] [32,33]. However, to our knowledge, there has been no explicit attempt to evaluate the costs and benefits of extracellular enzyme production in the context of the detailed metabolic budget of a cell.

The methodology of stoichiometric modeling is a natural avenue for addressing this challenge. The flow of biochemical reactions in metabolic pathways, including those involving extracellular enzymes, may be viewed as a complex resource allocation problem [28,34,35], which can be solved mathematically at the genome scale using constraint-based stoichiometric approaches [36,37]. These methods include Flux Balance Analysis (FBA) [36–38], initially developed for estimating the metabolic activity of individual species, and recently extended to the study of microbial communities [39–42] [5,10,43]. Dynamic Flux Balance Analysis (dFBA) [44], which iteratively updates pools of extracellular metabolites based on the fluxes observed in sequential executions of FBA, has proven particularly useful towards this goal [9,45], including through a framework called COMETS (Computation of Microbial Ecosystems in Time and Space) [9,46], which further extends dFBA to simulate the spatio-temporal organization of microbial communities [9,47–51]. Despite encoding complete mechanistic models of an organism’s metabolic network, FBA and dFBA approaches typically lack an explicit representation of extracellular enzyme reactions. The implicit implementation of extracellular enzyme function currently adopted in standard FBA/dFBA has fundamental limitations (See Methods), that make it impossible to effectively account for these processes in ecosystem-level models.

Here we embed extracellular enzyme production and activity within a model of the complete metabolic network of a microbial cell, and use the ensuing framework to simulate growth on an externally degradable resource. We address the limitations of previous approaches by introducing a modified version of dFBA that can explicitly account for the secretion and activity of extracellular enzymes. Our implementation, developed for the dFBA algorithm that is part of COMETS, treats extracellular enzymes as costly secreted metabolites [52], whose associated reactions take place in the extracellular environment according to explicit kinetic rate laws. We illustrate these new capabilities by studying an engineered yeast strain which constitutively expresses an exogenous cellulase [2]. In addition to displaying and testing our new COMETS module, we study the balance of costs and benefits in this engineered yeast system. We test the hypothesis that an optimal enzyme production strategy exists, which maximizes the net benefit to the organism, and study the dependency of this optimum on cellular and environmental parameters.

## Results

### A dynamic flux balance framework for extracellular enzyme secretion and function

We implemented, within the COMETS framework [46], a computational infrastructure to model extracellular enzyme activity in spatially structured microbial populations and communities. In making this modification to the basic structure of dFBA within COMETS, we connect the extracellular enzyme production and secretion to internal resource allocation in the cell. While the details of the implementation are described in detail in the Methods section, we provide here an overview of the approach, which is in itself novel and broadly generalizable.

Our formulation adds enzyme production and secretion as a set of reactions in the existing metabolic network (Fig. 1). These reactions, similar to the biomass producing reaction, transform a set of amino acids into an enzyme protein. The enzyme is then transferred into the extracellular environment, in analogy to other metabolic byproducts. Once an enzyme molecule has been secreted, it will be subject to physical and chemical processes equivalent to those that may act on other extracellular metabolites when applicable, namely diffusion and, potentially, degradation. A key parameter, which we name α, defines the amount of extracellular enzyme produced relative to biomass growth in units of millimoles per gram of dry weight (Fig. 1, Eq. 4). For example, if α = 0.001 mmol/gDW, for every gram of biomass produced, 0.001 mmol of enzyme will be secreted into the extracellular medium (for a protein with a molar mass of 142.1 kDa as used here, the secreted protein would represent 12.4% of the net biomass produced). This is the main parameter that defines the secretion strategy of the cell. While, as discussed later (see Discussion) one could implement α values that dynamically depend on environmental conditions, we are assuming here that α is a fixed parameter.

**Figure 1:**
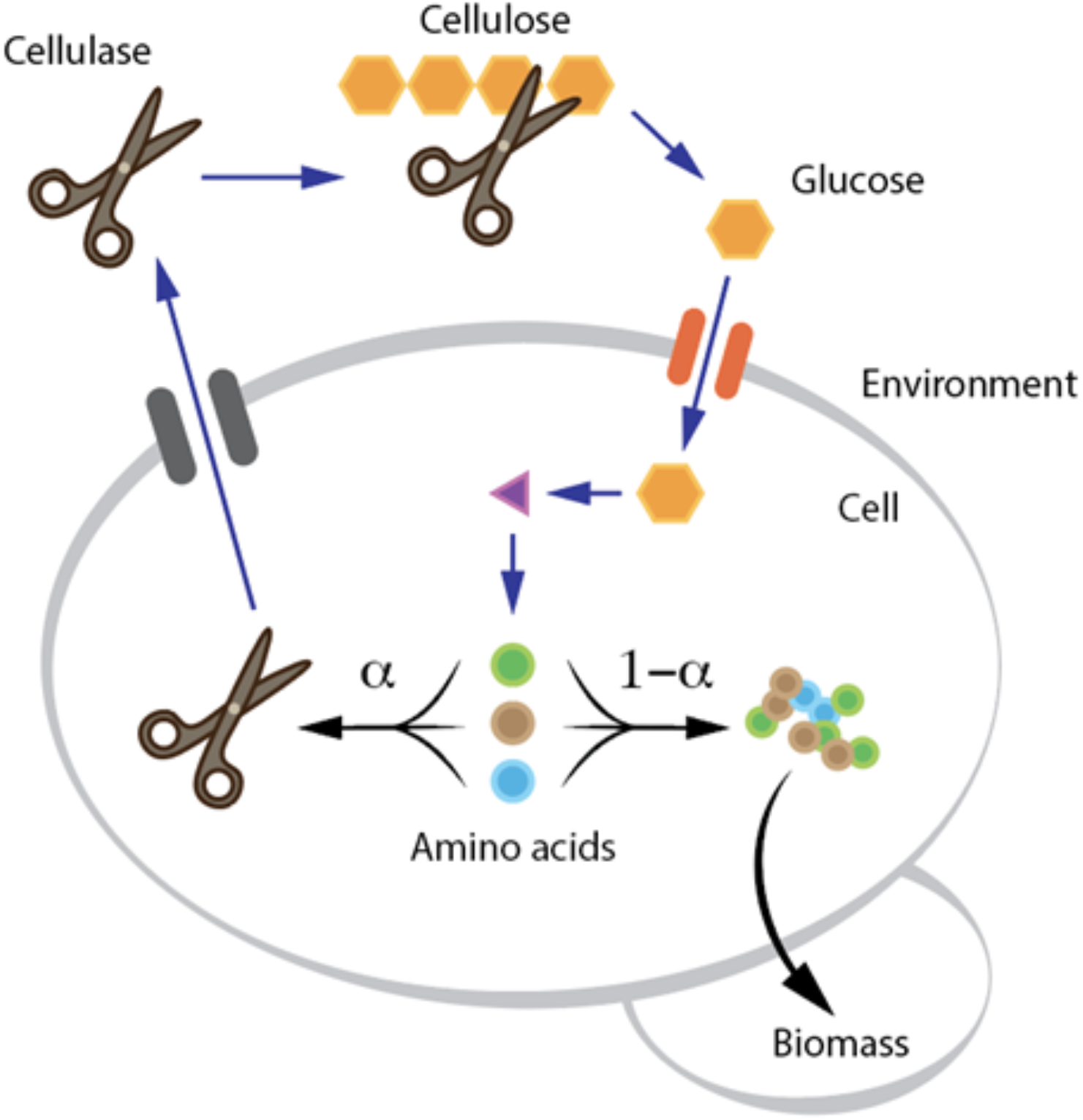
Simplified map of the reactions involved in cellulase secretion and cellulolysis, highlighting the key features of the system. The metabolic cost of cellulase production is taken into account by modeling its biosynthesis as a reaction that draws amino acids from the same pool used for biomass generation. Secretion of the enzyme into the environment is made proportional to growth by coupling the enzyme transport flux to biomass production. The enzyme present in the medium catalyzes the degradation of cellulose into glucose through a classical Michaelis-Menten mechanism. Notably, as opposed to previous implementation of cellulolysis in FBA, cellulose degradation can happen extracellularly independently of the organism that secreted the enzyme.

The next step we have taken to incorporate extracellular enzymes in COMETS is the explicit modeling of enzymatic activity in the extracellular environment. In this instance we have simplified the process by assuming the direct conversion of cellulose into glucose monomers, though intermediate reactions and metabolites can also be supported. Outside of the cell, a secreted enzyme is able to diffuse and perform its function independently of the organism by which it was produced. In other words, the enzyme becomes a common good that could catalyze reactions anywhere in the environment, and benefit any organism that could use its products. The central component of the simulation that makes this possible is the implementation of Michaelis-Menten kinetics for the enzyme in the extracellular environment: at every time step, in addition to performing FBA and diffusion, COMETS checks the presence of extracellular enzymes and corresponding substrate, and estimates the amount of substrate consumed and product(s) produced, based on enzyme-specific K_M_ and V_max_ parameters.

In our new module, extracellular enzymes are also subject to decay, e.g. due to thermal instability or proteolysis [53,54][54–56]. The simulation of enzyme degradation implies that the effort an organism puts into secreting an enzyme has a reward that decreases over time, making it impossible, for example, for an organism to benefit for an indefinite amount of time from the catalytic activity of an individual enzyme protein. Extracellular enzyme degradation is also important ecologically, as it can contribute to the generation of dissolved organic matter, whose decay products may be reused by other organisms or be part of broader biogeochemical cycles, particularly the cycling of nitrogen [57]. While the explicit generation of decay products is not implemented in the current version of COMETS, it would constitute a feasible addition.

### Implementation and testing of COMETS simulations for a cellulase-producing yeast

For the purpose of testing the new capabilities and exploring their potential to provide biological insight, we chose to focus on a modified *Saccharomyces cerevisiae* yeast strain, previously engineered to produce extracellular cellulases [58]. This strain is also part of an effort to implement consolidated bioprocessing, e.g. for bioethanol production from plant biomass [59,60] (See Methods for more details about strain choice and corresponding stoichiometric reconstruction). Using the engineered yeast strain (based on a modified version of the Yeast8 reconstruction, see Methods) and the newly updated COMETS framework, we simulated growth of this yeast in a homogenous anaerobic environment with a limited amount of cellulose as the main carbon source and an excess of all other essential metabolites (see Methods, and Supplementary Table S.4).

We found that our new implementation of extracellular enzyme activity in COMETS was able to recapitulate the in-vitro growth curve of cellulase-expressing *S. cerevisiae* (Figure 2). Simulated growth best matched experimental data at intermediate values of α (α = 0.0032 mmol/g), using kinetic parameters of the extracellular enzymes reported in the literature (V_max_ = 32 s^−1^ and K_M_ = 0.2 mg/mL [61,62]). This value of α would represent the expected extracellular enzyme secretion rate for the simulated *S. cerevisiae* strain. Interestingly, this value of α also falls within the range of cellulase production measured in this engineered yeast for endoglucanase(α = 0.0026) and β-Glucosidase (α = 0.0072) (see [63] and Methods).

**Figure 2:**
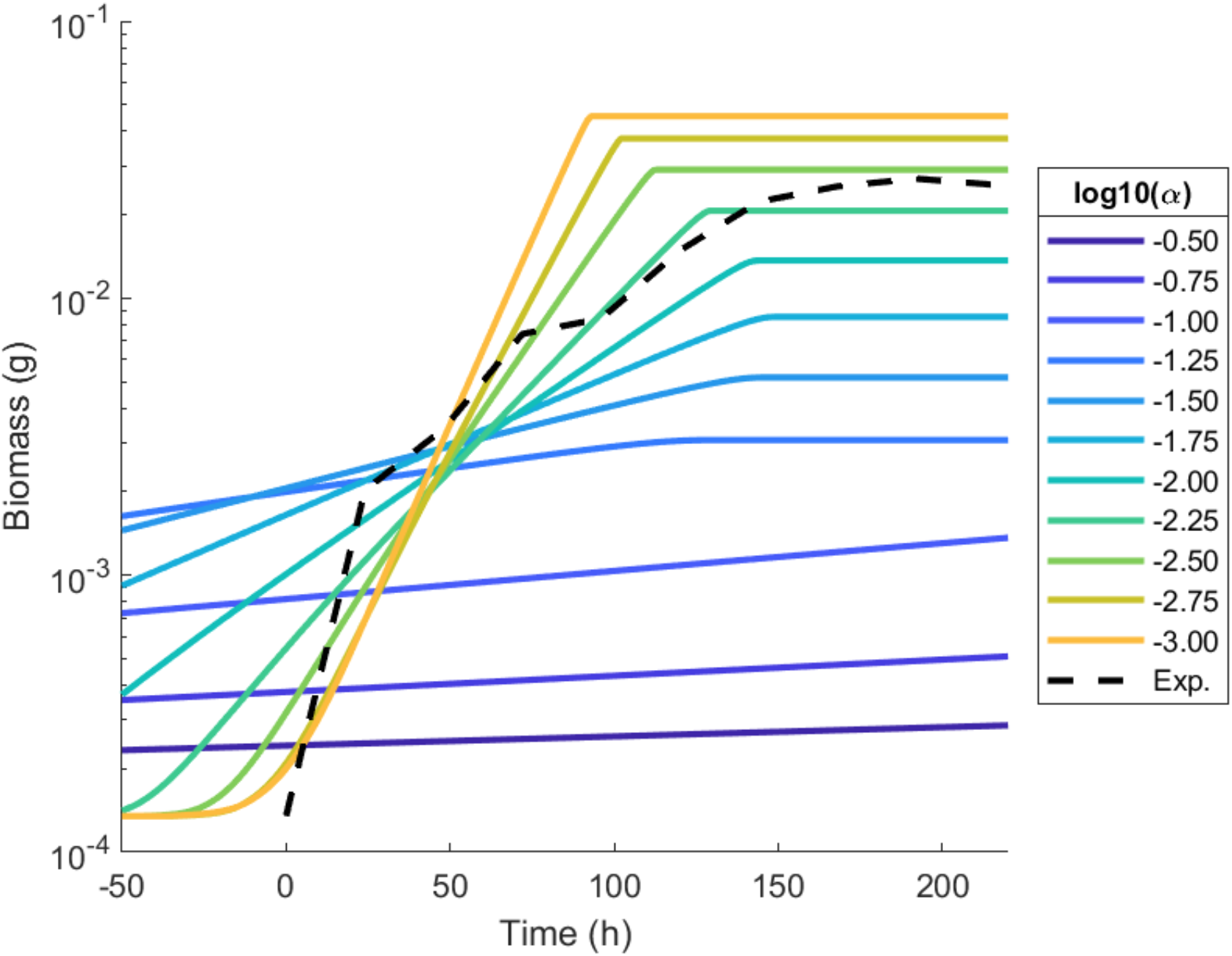
Comparison of the in-vitro growth (dotted line, data from [2]) of cellulase-expressing *S. cerevisiae* to simulations using a range of values for the cellulase expression rate α. Each simulation data series has been time-shifted to the values which provide for the closest fit to the experimental data from [2]. The best fit was achieved with α = 0.0032 mmol/gDW, with an RMSE of 0.2375.

We found that a similar value of α is found even if we simultaneously search for α and enzymatic parameters that best fit the experimental curve. The values of the parameters obtained by fitting to a linear regression which allowed the simulation to most closely match the *in vivo* data were V_max_ = 87.5 s^−1^, K_M_ = 0.0094 mg/mL, with α again at a value of 0.0032 mmol/gDW, with an RMSE of 0.1659. It was reassuring to see that the kinetic parameters are within an order of magnitude of the expected values from the literature [61,62] and that the ideal α remains unchanged under both optimization procedures, suggesting that its estimate is robust relative to uncertainty in other parameters. This value of α is also consistent with previously reported rates of cellulase production [64–66]. The literature-derived enzymatic parameters are used as the defaults for all subsequent simulations in this report, unless otherwise noted.

### Balance between cost and benefit points to resource-dependent optima

Different values of α strongly influence the dynamics of the system (Figure 3). In particular, the trajectories of glucose and cellulose concentrations and the change in biomass over time vary not only quantitatively (e.g. initial rates) but also qualitatively as a function of the rate of enzyme secretion. If the value of α is too small, there is virtually no cellulose to glucose conversion (Fig. 3A), and hence very little growth. Conversely, if α is too large, cellulose is converted to glucose very quickly (Fig. 3B); however, the expense for extracellular enzyme production leaves few resources available for biomass production, making growth asymptotically slow (Fig. 3D and E). This output supports our hypothesis that there should be a balanced enzyme production strategy which maximizes the utility of extracellular enzymes [28–31]. The existence of an optimum was confirmed by systematically repeating simulations for many values of α, and plotting the corresponding growth rate and yield (Fig. 4).

**Figure 3:**
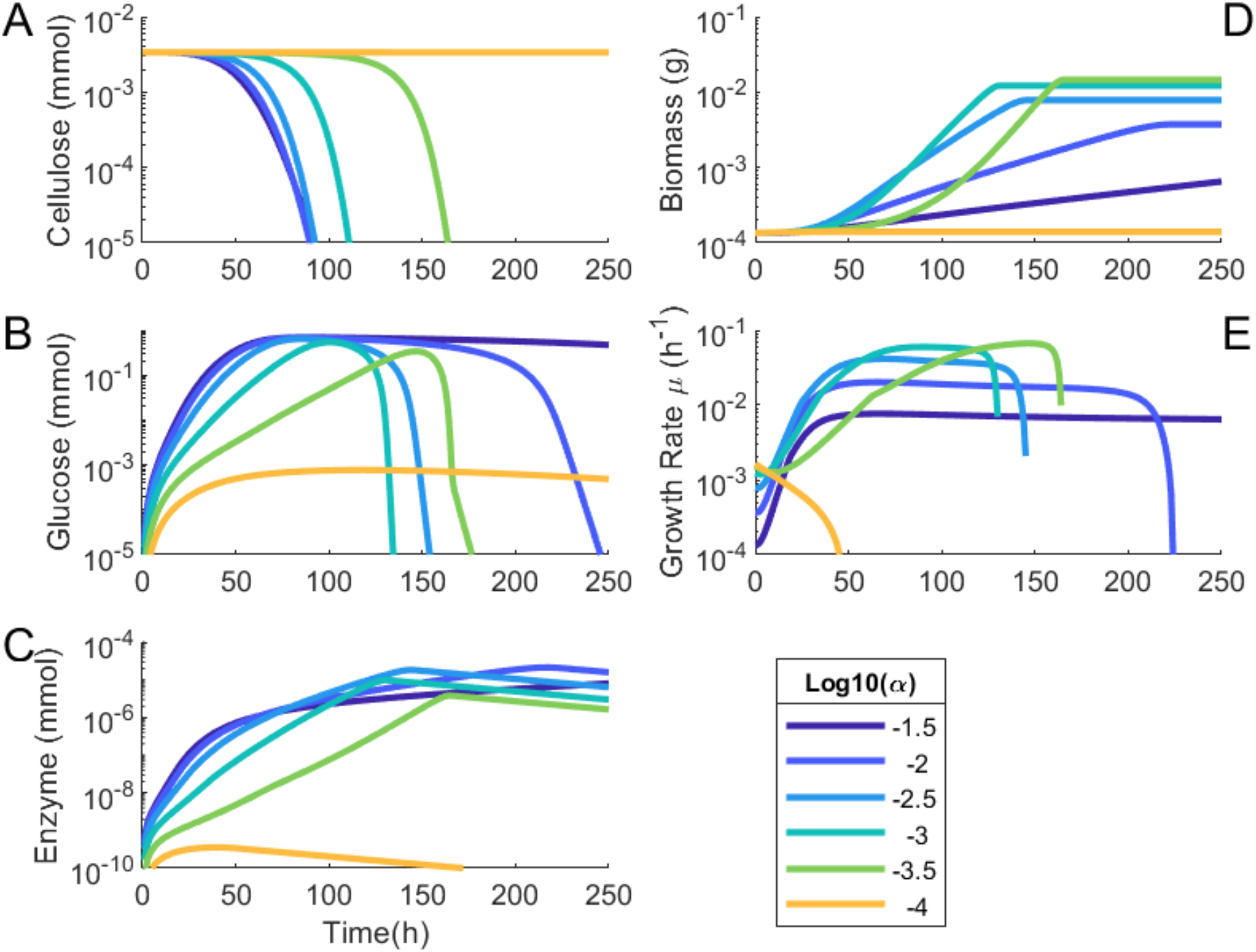
Logarithmic plots of cellulose (A), glucose (B), and enzyme (C) concentrations, biomass (D), and growth rate (E) over time for simulations using different enzyme production rates. A) Cellulose degrades over time in proportion to the amount of enzyme in the system. B) Glucose accumulates when cellulolysis outpaces the capacity of the yeast to consume it, and this accumulation is faster in systems with higher enzyme secretion rates. C) The amount of extracellular enzyme increases in proportion to biomass growth. D) Total biomass and E) growth rate increase over time with the greatest increases seen in the models with intermediate α values.

**Figure 4:**
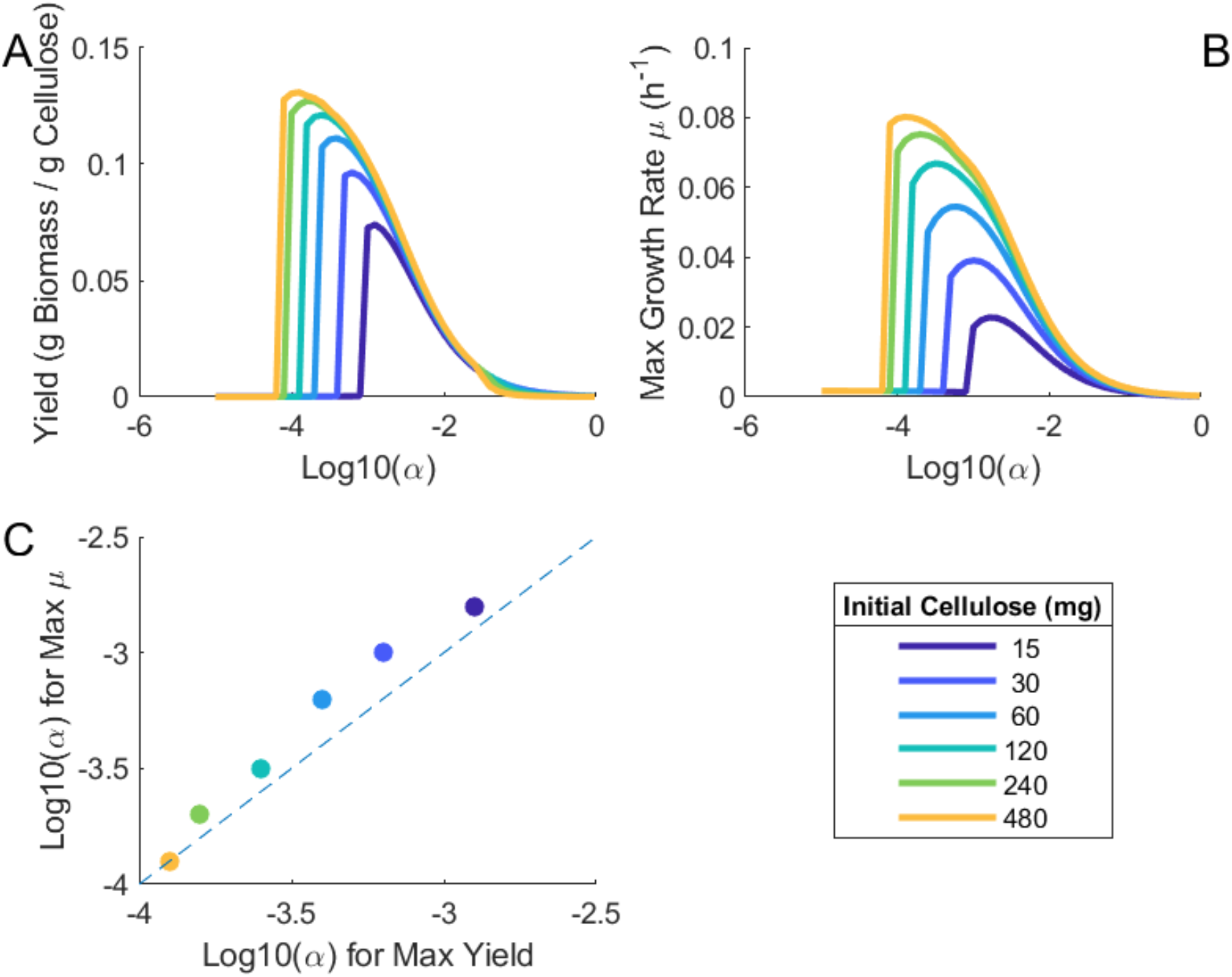
Yield (A) and maximum growth rate (B) in homogenous simulations with varying amounts of cellulose reveal that optimal α decreases for environments with increased cellulose. (C) Comparison of α values associated with maximum yield (x axis) and rate (y axis) for different initial amounts of cellulose. The graph displays a strong positive correlation, indicating that enzyme production rates that maximize rate are predicted to also maximize yield.

The α at which this optimum is achieved, and the corresponding maximal values of yield and rate (which are correlated to each other, Fig. 4C), depend strongly on the initial cellulose available. Curves for growth and yield as a function of α have a characteristic asymmetric shape (Figs 4A,B), with a sharp decline at the low end of the α axis. This sharp decline happens very close to the optimal α, implying that the ideal cellulase production rate is critically close to a value at which the organism does not produce enough enzyme to guarantee growth. Production of excessive amounts of enzyme will result in a lower yield and growth rate but does not pose the same risk. This predicted shape of the curve has a couple of possible consequences: first, it implies that it might be safer for an organism to overshoot with a larger value of α, and gradually reduce it (through physiological or evolutionary adaptation), rather than approaching the optimal value from below. Second, it suggests that the pressure of natural selection on constitutive enzyme production favors underestimating the abundance of resources in the environment.

## Discussion

Despite the broad importance of extracellular enzyme processes across environmental conditions, mainstream efforts to model genome-scale and ecosystem-level metabolism have so far circumvented the problem of how to appropriately embed these effects in constraint-based models. We developed a framework to explicitly simulate extracellular enzyme functions in dynamic FBA in a way that is simple enough to enable a systematic analysis of the role played by different key parameters, but realistic enough to allow a qualitative comparison with experimental systems. By applying this new framework to a specific case study, we were able to computationally test the hypothesis that a quantifiable optimal allocation of resources to extracellular enzymes exists as a function of substrate availability.

The integration of extracellular enzyme secretion and function in COMETS is an important step towards understanding the ecology and evolution of microbial community dynamics in environments rich in complex macromolecules. Our simulations suggest the existence of optimal enzyme production rates relative to biomass production in a way that depends on the environmental availability of substrate. Importantly, this finding is the complex outcome of several factors that would not be available in simple partial differential equations models of microbial growth, or in standard FBA or dFBA simulations. The existence and specific location of these optima depends on the properties of the intracellular metabolic circuits of the organism (e.g. the value of α), the extracellular enzyme parameters, and the complex time-dependent interplay between the different quantities (cellulose left, glucose generated, biomass present, etc.). While the organism, in a dFBA simulation, has an instantaneous objective function (as in regular FBA), the potential optimal behavior under a given environment is not captured by that objective, but requires taking into account these full complex dynamics. Growth optima can strongly depend on the ecological setting. In a competitive environment where space or resources are scarce, the enzyme produced by one organism can become a precious common good (together with the cellulolytic byproducts themselves), enabling cheater organisms to benefit. However, in an environment where the common good is shared with an organism that is producing other useful resources, mutualistic relationships may enable diverse communities to thrive [67–69]. Our analysis provides an expected baseline of how specific rates of extracellular cellulase secretions would affect fitness of an organism in isolation. However, the flexibility of COMETS and the availability of detailed genome-scale models for many organisms for different biomes will enable detailed in silico experiments with multi-species communities, where the rise of cheaters and the complexity of multi-species competition and exchange may modulate the optimal choice of enzyme secretion.

A number of decisions were made in the design of this study to keep the modeled system as simple as possible, while remaining biologically informative. One simplifying assumption, which limits the current reliability of specific quantitative predictions, is the implementation of cellulolysis as a single-step process. Cellulolysis in natural systems typically involves at least three classes of enzymes (endocellulases, exocellulases, and beta-glucosides) responsible for different steps of the conversion of cellulose into simple sugars. Each of these reactions has distinct kinetic parameters that may be fine-tuned. These pathways display significant synergistic effects both within and across classes of cellulases [70]. Here we approximate these processes as a single enzymatic reaction, converting cellulose directly into glucose. While it is reassuring that, despite this simplification, we could recapitulate some features of a specific experimental system, based on parameters from the literature, it will be important, in future developments, to explicitly introduce the individual enzymes, and corresponding steps. This is already feasible with our new COMETS module for extracellular enzyme secretion, but it will require introducing individual α values for each enzyme. Moreover, while current simulations were using a pure and relatively simple cellulose substrate (Avicel), simulations of plant biomass degradation by microorganisms in natural environments will have to deal with much more complex mixtures of substrates, including lignins and hemicelluloses. In addition to explicitly modeling the corresponding degradation pathways, simulations including these factors could be implemented by introducing simplified phenomenological rules that represent the conversion of inaccessible cellulose into an accessible form, potentially taking advantage of the spatial features of COMETS.

Future efforts could build upon our current approach in a number of directions. First, while the system modeled here is a genetically engineered yeast which constitutively secretes its extracellular enzymes, it will be interesting to expand our current framework to introduce regulatory control and different modes of enzyme production and secretion in response to environmental stimuli. We expect that even simple feedback systems, e.g. based on sensing of sugars will be able to significantly reduce the metabolic burden of the enzyme expression system. The implementation of similar sensing and regulatory mechanisms would constitute a broadly interesting research avenue in extensions of dFBA simulations. For example, one could include product inhibition or treat the enzyme secretion rate as a nonlinear function of the growth rate (in contrast to the direct proportionality used here). Metabolic models can even be created that switch between multiple strategies based on environmental conditions. Second, our work paves the road for addressing a number of questions related to the spatial distribution of biomass, enzymes, and carbon sources, which can be simulated using the spatial features of COMETS. The spatial component would be particularly important for addressing questions about the evolution of cooperation and competition induced by the diffusion of extracellular enzymes and their byproducts in multi-species communities. Finally, while we have presented here a limited sample of the capabilities enabled by our extension to COMETS, this approach is easily extensible to other natural processes. We have focused on a simplified cellulolysis system, but other degradation enzymes, such as proteases or alginases, can be implemented directly with the enzyme reaction method described here. Metabolite-sequestering molecules like siderophores can be modeled through the addition of binding and unbinding reactions. Thus, the extension of dFBA and COMETS proposed here can find useful applications in multiple areas of metabolic engineering and microbial ecology, including studies of soil and plant-associated communities, the human microbiome, marine microbial communities, and the production of pharmaceutical and commercial products in bioreactors. Furthermore, we should note that the COMETS module that implements extracellular reactions is very general, and allows users to define any arbitrary number of chemical reactions happening extracellularly with or without the presence of an enzyme. This feature could have a number of additional applications, including for the study of early biochemical evolution [71].

## 5 Methods

### Dynamic Flux Balance Analysis and COMETS

Flux balance analysis (FBA) is a well established approach for using stoichiometry, a steady state approximation and optimality principles to predict genome-scale metabolic fluxes. It has been described extensively elsewhere [36–38]. Its extension to the temporal domain, dynamic FBA (dFBA), still assumes that intracellular metabolites quickly equilibrate to a steady state, but explicitly simulates changes in the biomass abundance of different species, and in the molecular composition of the environment [9,44,72]. This aspect of dFBA makes it appropriate as a platform for modeling extracellular metabolic processes using standard chemical kinetics to allow molecules in the media to interact without any direct dependence on microbial biomass or metabolic models. Within the COMETS (Computation of Microbial Ecosystems in Time and Space) framework [9,46,73], dFBA has been embedded in a spatial environment, where metabolic processes are accompanied by the physical processes of molecular diffusion and biomass expansion upon growth. COMETS can therefore be used to study the spatio-temporal organization of microbial communities, with applications in multiple areas of microbial ecology [9,47–51].

Current FBA and dFBA formulations can incorporate a description of extracellular metabolic activity, but with major limitations, which constitute part of the motivation for the present work. In traditional FBA, for example, the conversion of Cellulose (C) to Glucose (G) by an extracellular (membrane bound) cellulase would be represented as follows:

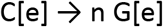

where [e] symbolizes the extracellular compartment, and n is a hypothesized number of glucose monomers obtained by cleavage of one cellulose polymer. This reaction would be part of the metabolic model of the organism that produces the cellulase.

Under this standard model’s assumption, the enzymes associated with extracellular reactions cannot be decoupled from the model, and would therefore disappear if the organism was removed [74]. Furthermore, by handling these catalytic functions in an organism-dependent way, the existing approach constrains them to the immediate vicinity of the cell. This approach also creates unrealistic estimates of enzyme abundances and rates, e.g. by preventing the depletion of enzymes over time or their accumulation at a rate different from that of growth. Finally, the classic approach does not take into account the cost of producing enzymes that are effectively lost biomolecules from the perspective of the overall cellular metabolic budget.

### Addition of extracellular reactions in COMETS

We have included in COMETS the capacity to handle two types of reactions which can take place in the extracellular environment. The setup for these two types of reactions is instructed in the COMETS simulation layout definition file (see [46,73]). The first type are simple mass-balance equations. A set of reactants (with a given stoichiometry) react at a given rate *k* to produce a set of products (also with a given stoichiometry):

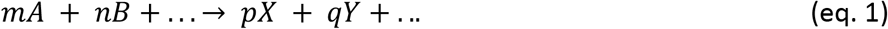

This would translate into the following differential equation for the production of one of the reaction products (for example X), which COMETS uses to estimate the change of X during the next time step:

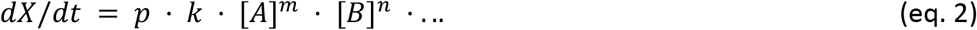

This formula can also be applied to model other processes of arbitrary order, such as metabolite decay or addition of media in a chemostat. In particular, in this work, we use this process to simulate enzyme (E) degradation in the extracellular environment. In this case, the reaction can be written as a special case of eq. 1 above, i.e. E → Null. We should point out that this is effectively an exponential decay (E(t) = E_o_e^−*k*t^) with decay rate *k*, provided as an input by the user.

The second class of reactions that have been implemented are monomolecular enzyme-catalyzed reactions. Enzymatic reactions follow the Michaelis-Menten equation for rate *v* = *V_max_* · *E* · (*S* / [*K*_*M*_ + *S*]), with enzyme and substrate concentrations *E* and *S*, the turnover rate *V_max_*, and the half saturation constant *K_M_*. As with defining mass-balance reactions, reactants and products are defined with a fixed stoichiometry by the user. A single extracellular metabolite is defined as the enzyme which carries out the reaction, and is given values for the catalytic rate (*V_max_*) (units: s^−1^) and Michaelis constant (*K_M_*) (units: mM). The concentration of the enzyme in the media is not altered by its participation in an enzymatic reaction.

The dFBA simulation process in COMETS involves cycles comprising several steps. The first step in each cycle (updating metabolite concentrations based on cellular reaction fluxes) is unchanged relative to standard dFBA [9]. We have now added an extra step, right after this first step, in which COMETS handles extracellular reactions. COMETS converts the input set of extracellular reactions as defined by the user into a system of differential equations and approximates a solution with the Runge-Kutta algorithm (RK4) implemented via the Apache Commons Math API version 3.4 for Java [75]. In instances where reactions occur fast enough that the algorithm predicts a negative concentration of any metabolites at the end of a single timestep, the current timestep is recursively subdivided into two consecutive reaction steps. Each coordinate cell in the simulated space is run through the ODE solver using its initial metabolite concentrations as the starting conditions, and the result becomes its new set of concentrations.

The kinetics of the extracellular reactions was validated in COMETS by implementing simple simulations which do not involve the FBA component, only the reaction, enzymes, and substrates. These simulations adhere to predicted rates for both exponential decay and Michaelis-Menten reactions (Fig. S1).

### Enzyme Production and Secretion in a Stoichiometric Model

The reaction for the production of an extracellular enzyme from precursor amino acids is added to the metabolic model in the form:

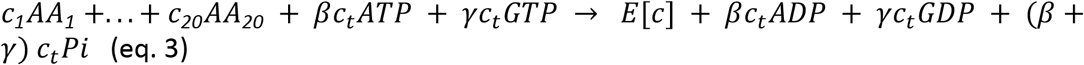

where *c_i_* is the number of amino acid *i* that occurs in the protein’s sequence (Table S4), *c_t_* is the total length of the polypeptide sequence, and β and γ are constants which define the energy cost of polymerization (Table A.1). This reaction effectively produces the enzyme as if it was an intracellular metabolite. As shown next, however, the only fate of this enzyme is extracellular secretion.

We assume here that the rate of secretion of the enzyme is proportional to the growth rate. This coupling between growth rate and enzyme secretion could be achieved in multiple ways (including in principle creating a new biomass production reaction that also secretes the enzyme). In our current implementation, we achieved growth-coupled enzyme secretion by adding a fictitious “Coupling Metabolite” (CoupleMet) as an additional product of the biomass reaction. This Coupling Metabolite is then coupled to the transport of the enzyme from the cytoplasmic ([c]) to the extracellular ([e]) environment. Thus the overall biomass-coupled enzyme secretion is encoded in the two following reactions:

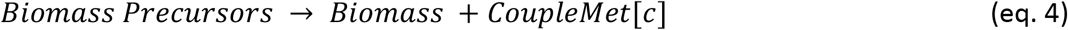

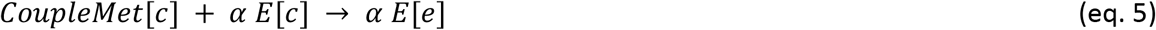

The parameter *α* represents the amount of enzyme produced per unit of biomass produced, and is one of the key parameters explored throughout the paper, as it captures the cellular choice of how much resource investment the cell should put into producing the enzyme vs. growing. Note also that once out of the cell, the extracellular enzyme E[e] behaves like a metabolite, except for its capacity to catalyze the associated extracellular reaction.

### Initial conditions for bootstrapping growth

It is important to note that the carbon used by the metabolic model for the majority of growth and fermentation in this case study is only available in the form of glucose, generated as the product of enzymatic cellulolysis. Therefore at the beginning of the simulation, unless a usable carbon source is provided (in addition to cellulose), cells would not be able to bootstrap themselves into growth. In the current simulations, we thus provide a small amount of mannose to initialize growth and enzyme production. This mannose is quickly consumed, leaving the yeast to grow solely on glucose produced by cellulose degradation. We provide mannose, as opposed to glucose, in order to be able to easily track the glucose produced as a product of cellulose degradation. Alternative strategies for bootstrapping initial growth would be feasible. These include the assumption of an initial amount of intracellular carbon storage or an initial amount of extracellular enzyme in the environment. Simple tests we implemented with these alternative approaches provided growth curves substantially equivalent to the ones shown, and were not pursued further. Notably, it is important to design the bootstrapping phase as a short transient relative to the experiment itself. Here we chose an initial mannose amount of 0.01598 mmol. Changes in this initial amount would be roughly equivalent to changes in the initial amount of biomass and enzyme (see sensitivity analysis in supplementary data).

### A cellulase-producing yeast as a model system

The modified cellulase-producing *S. cerevisiae* system has several advantages that make it appropriate for our study: (i) It can take advantage of a well established high-quality stoichiometric reconstruction, i.e. the *S. cerevisiae* stoichiometric model, which has a long history of manual curation and validation [76,77]; (ii) Yeasts serve as a common chassis for genetic engineering [78–80]; (iii) This particular system is associated with a clear well-studied laboratory equivalent, for which experimental data are available; Growth data and cellulose degradation dynamics were taken from an experiment performed by Den Haan et al. [2] utilizing a *S. cerevisiae* strain genetically modified to constitutively express the exogenous cellulase genes *Saccharomycopsis fibuligera* BGL1 and *Trichoderma reesei* EGI; (iv) Fungal degradation of plant biomass is highly relevant both for metabolic engineering and ecological applications, making it a system which can be expanded further, with broad implications. It uses enzymes from widespread plant decomposing fungi, and can serve as a valuable proxy for obtaining insight relevant for important open ecological questions.

In order to recapitulate the above experimental system, we imported in COMETS the most recent stoichiometric reconstruction for *S. cerevisiae* (Yeast8 [76]). To construct a model of the specific engineered strain used in [2], we modified the anaerobic Yeast8 yeast metabolic model so as to make it able to produce and secrete a cellulase enzyme, as described above. Note that while the original modified organism was engineered to secrete both endoglucanase and beta-glucosidase, we implemented a simplified version in which a single representative cellulase enzyme is secreted. This cellulase is then assumed to degrade cellulose into glucose in a single-step reaction, in analogy with previous implementations in stoichiometric models [81,82]. The extracellular cellulolysis reaction was added to the simulation with the following parameters: 218.2 units of glucose were produced for each unit of cellulose consumed, in accordance with the average degree of polymerization of Avicel [83]. Initial estimates for turnover rate V_max_ and half-saturation constant K_M_ were 32 s^−1^ and 0.2 mg/mL respectively, based on values reported in BRENDA for BGL1 [84].

Note also that genome-scale metabolic models typically include a growth-associated (GAM) and a non-growth-associated maintenance (NGAM) flux, which are set to fixed experimentally determined values, to take into account cellular metabolic demands not directly associated with metabolism itself. For the current simulations, in order to simulate some degree of internal metabolic activity also in absence of growth, we used a NGAM of 0.7 mmol/gDW·h (same as anaerobic growth, as in the reported aerobic case [85,86]). In parallel we adjusted the GAM from 30.49 mmol/gDW·h to 23.49 mmol/gDW·h in order to match experimental growth rates. In any case, supplemental figure S4 demonstrates that the system is robust to a wide range of values for these parameters.

### COMETS Simulations

Simulations for all experiments were performed with COMETS version 2.9.0 [46] utilizing Gurobi version 9.0 [87]. Input file generation and output processing were performed using the COMETS Toolbox for Matlab [46]. Unless otherwise indicated, experiment parameters were set to recreate the batch culture conditions in [2]. Culture volume was 100mL. Initial cellulose amount was set to a 0.0125 millimolar concentration (equivalent to 2g Avicel in 450mL stock), and all other essential metabolites were set to 100 millimolar concentrations so that they would not be limiting factors. Enzyme decay was set as a first-order reaction [56] eliminating the enzyme at a rate of 1% per hour. Simulation time step length was set to 1 hour. Cell death was disabled in this simulation. K_M_ for glucose uptake by the organism was set to 3.94 mmol/cm^3^, and the maximum glucose uptake rate was set to 5.6e-4 mmol/gDW*h. The initialization of growth in these simulations requires inclusion of a carbon source that can be utilized by the metabolic model. This can be achieved directly by including sugars in the initial media, or including a small amount of enzyme in order to begin the conversion of cellulose to glucose. Mannose was selected to be included in the media in order to keep the initial carbon source distinct from the glucose produced by cellulolysis. Conceptually, this initial source serves the same purpose as a yeast cell’s internal carbon stores, the depletion of which are not supported by a steady-state model. For the simulations shown here, in order to bootstrap initial growth before cellulolysis has begun, 0.015 mmol of mannose was added to the media and mannose uptake was enabled in the model with kinetics equal to those of glucose uptake (V_max_ = 3.9 mmol/gDW·h, K_M_ = 0.56 mM).

Fitting to experimental data was performed by a hill-climbing algorithm was used to tune these three parameters, determining the goodness of fit based on minimizing the root mean square error (RMSE) between the log_10_ of the the time-shifted simulated biomass and the biomass recorded in vivo at the reported times (see Supplemental Information for implementation details). Because the RMSE landscape is convex (Fig S3) a naïve hill-climbing method is expected to reach the global optimum for finite values of V_max_, K_M_, and α.

This is a simplification of the actual cellulolysis process, which is typically a multistep reaction involving three separate enzymes. Future work should examine the impact of this, but for now this approach is used in the interest of keeping the number of simulation parameters at a manageable size and to avoid overcomplicating the discussion of results by introducing intermediate metabolites.

### Inference of a range for α from experimental data

Data available from van Rensburg et al. [63] allowed us to roughly estimate a realistic range of values for the α parameter we define and use in our model. The estimation reported below is based on Fig. 1 and Table 4 from [63]. In particular, the table reports an estimate of endoglucanase produced of 0.013 (±0.001) mg protein ml^−1^. Given that the final amount of biomass in the bioreactor (as shown in Fig. 1) is approximately 5g/l, the proportion of extracellular protein produced per unit biomass is roughly α^Endo^ = (0.013 g/l) / (5 g/l) = 0.0026. A similar calculation for β-Glucosidase (produced in an amount of 0.036 (±0.015) mg protein ml^−1^) leads to α^βG^ = (0.036 g/l)/ (5g/l) = 0.0072. We thus infer the range 0.0026 - 0.0072 as a crude estimate of realistic values of α, to compare with our computational model optimization estimates (α=0.0032).

## Acknowledgements

We are grateful to members of the Segrè and Bhatnagar labs for helpful inputs and discussions at multiple stages of this work. All authors were supported by the National Science Foundation (Award 1457695) and the Boston University Interdisciplinary Biomedical Research Office. D.S. further acknowledges funding from the U.S. Department of Energy, Office of Science, Office of Biological & Environmental Research through the Microbial Community Analysis and Functional Evaluation in Soils Science Focus Area Program (m-CAFEs) under contract number DE-AC02-05CH11231 to Lawrence Berkeley National Laboratory; the NIH (T32GM100842, R01GM121950), the National Science Foundation (NSFOCE-BSF 1635070), the Human Frontiers Science Program (RGP0020/2016).

## Supplementary Material

**Figure S1:**
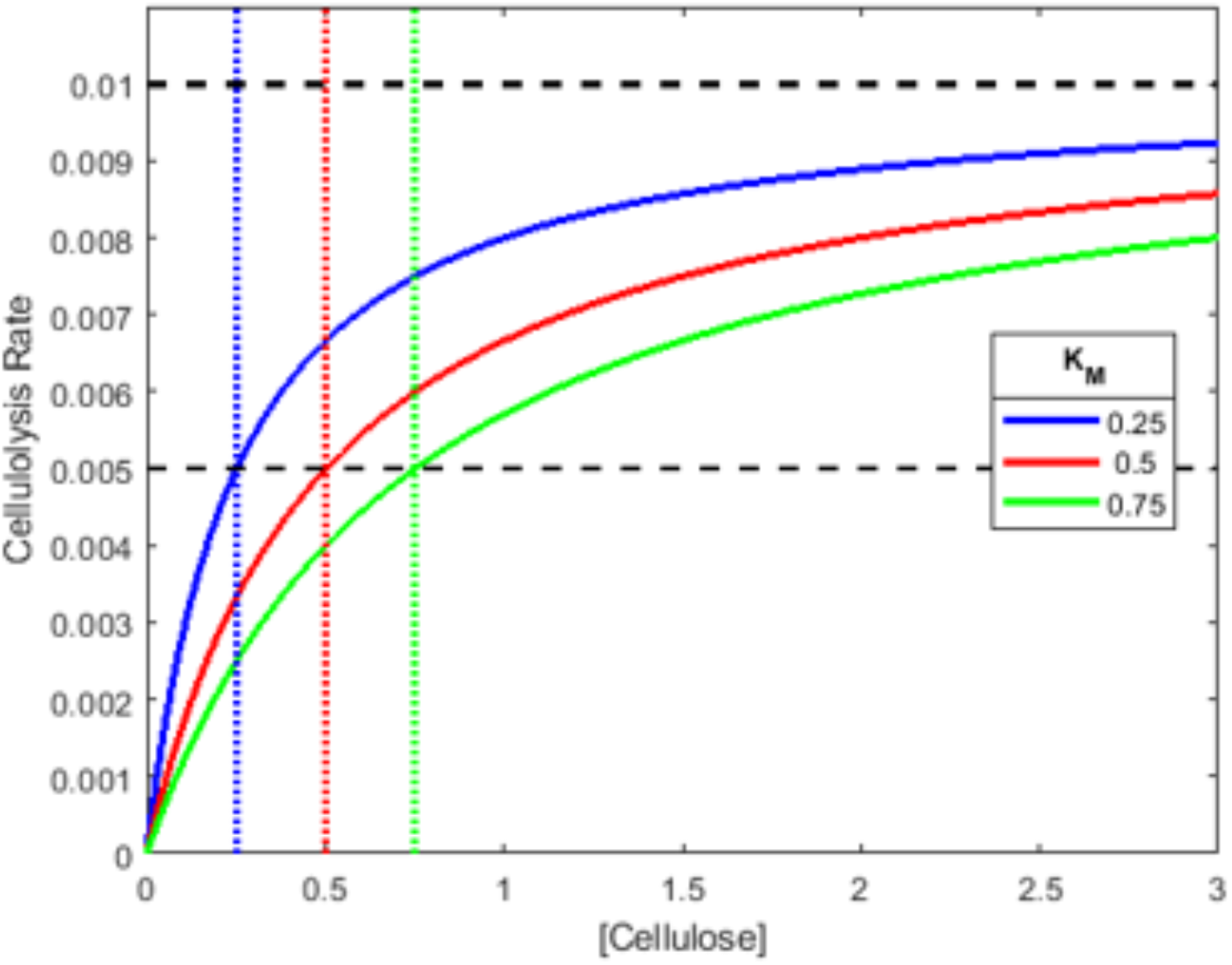
Extracellular enzymatic reactions in COMETS.

Enzyme-catalyzed extracellular reactions with a range of values for K_M_ adhere to the pattern expected of Michaelis-Menten reactions, namely achieving a rate equal to ½ V_max_ when substrate concentration is equal to K_M_, and asymptotically approaching V_max_ with increasing substrate concentration.

To confirm the behavior of enzymatic reactions, COMETS simulations were performed with an initial concentration of 3 mmol enzyme and 1 mmol substrate. An extracellular reaction was input for enzymatic degradation of the substrate following the Michaelis-Menten equation v = Vmax * [E] * [S]/(KM + [S]), for a Vmax of 0.01 s^−1^ and a range of values for K_M_. The rate of the reaction adhered to the pattern expected of Michaelis-Menten reactions, namely achieving a rate equal to 1/2 V_max_ when substrate concentration is equal to K_M_, and asymptotically approaching V_max_ with increasing substrate concentration (Fig S1). Exponential decay can be represented with an initial concentration of reactants in the media and no biomass. A first-order reaction was input for the decay of the metabolite with concentration N such that v = −λN for a range of λ = 0, 10, or 100 s^−1^. The decrease in a quantity N follows the law N(t) = N_0e_-t, which yields the reaction rate dN/dt = −N. A first-order reaction was input for the decay of the metabolite with concentration N such that v = −λN for values of 0, 10, or 100 s^−1^. Concentrations at measured timepoints exactly matched the expected values as calculated with the formula N(t) = N0e^−λt^ (series marked with X’s). Implementing this is simply a matter of defining an extracellular reaction which has a single reactant, X, a rate equal to −[N], and which yields no products (Fig S2).

**Figure S2:**
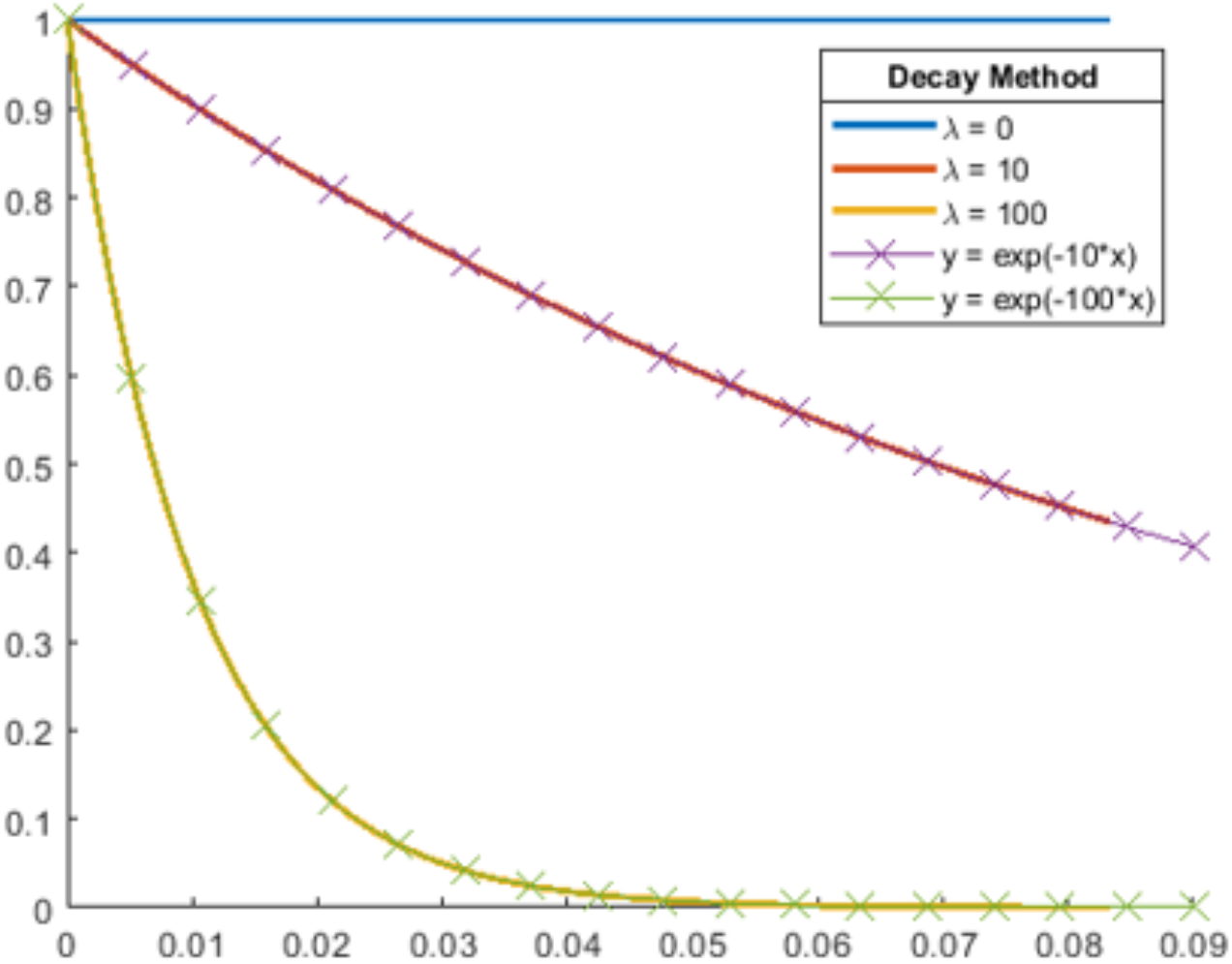
Fitting In Silico Growth to In Vivo Results.

Extracellular reaction behavior in COMETS was validated by running simulations with an initial concentration of reactants in the media and no biomass. Solid series depict time points measured in COMETS, while series marked with X’s show the predicted concentration according to the rate equation.

In order to fine-tune the kinetic properties of the cellulase enzyme and determine the enzyme secretion rate α which most closely adhered to a living system, in silico predictions for growth were prepared on a non-spatial simulation matching the conditions for an experiment carried out by Den Haan et al. (11). Initial cellulose was set to 0.54g to reflect the 27% enzymatic conversion of the 2g of Avicel used in vivo, and an excess of other essential metabolites was provided (oxygen, water, phosphate, sulfate, ammonium, potassium, and iron). The initial microbe population was set to 0.135 mg, and 0.015 mmol of mannose was added in order to initialize growth and enzyme production.

A simple hill-climbing algorithm was employed to find the best fit between the data while varying α, turnover rate, and the Michaelis constant in turn.

**Figure S3:**
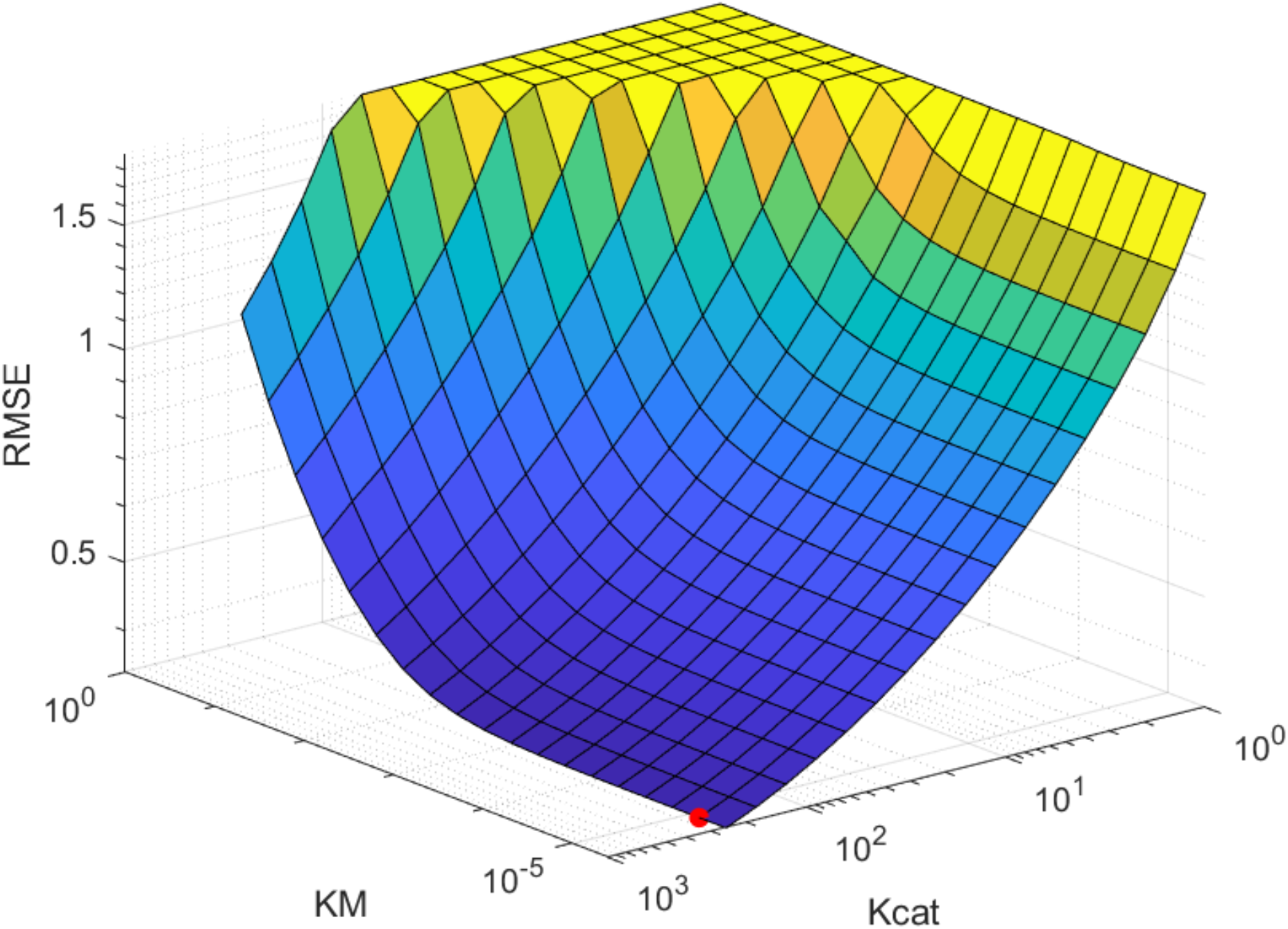
The Space for Fitting Enzyme Kinetics is Smooth.

The RMSE for the indicated parameter space over turnover rate and half-saturation constant as used to determine enzyme kinetics has a single valley. Red dot indicates the point where RMSE equals the minimum. Here α = 0.0032.

A simple approach for finding the k_cat_ and K_M_ constants was evaluated by graphing the root-mean-square error calculated via the difference between the observed and simulated growth curves over a range of each parameter. The error function is monotonic in regards to both variables, suggesting that a simple hill-climbing (or in this case, hill-descending) algorithm is sufficient to find parameters which match the behavior of the in-vivo system.

**Figure S4:**
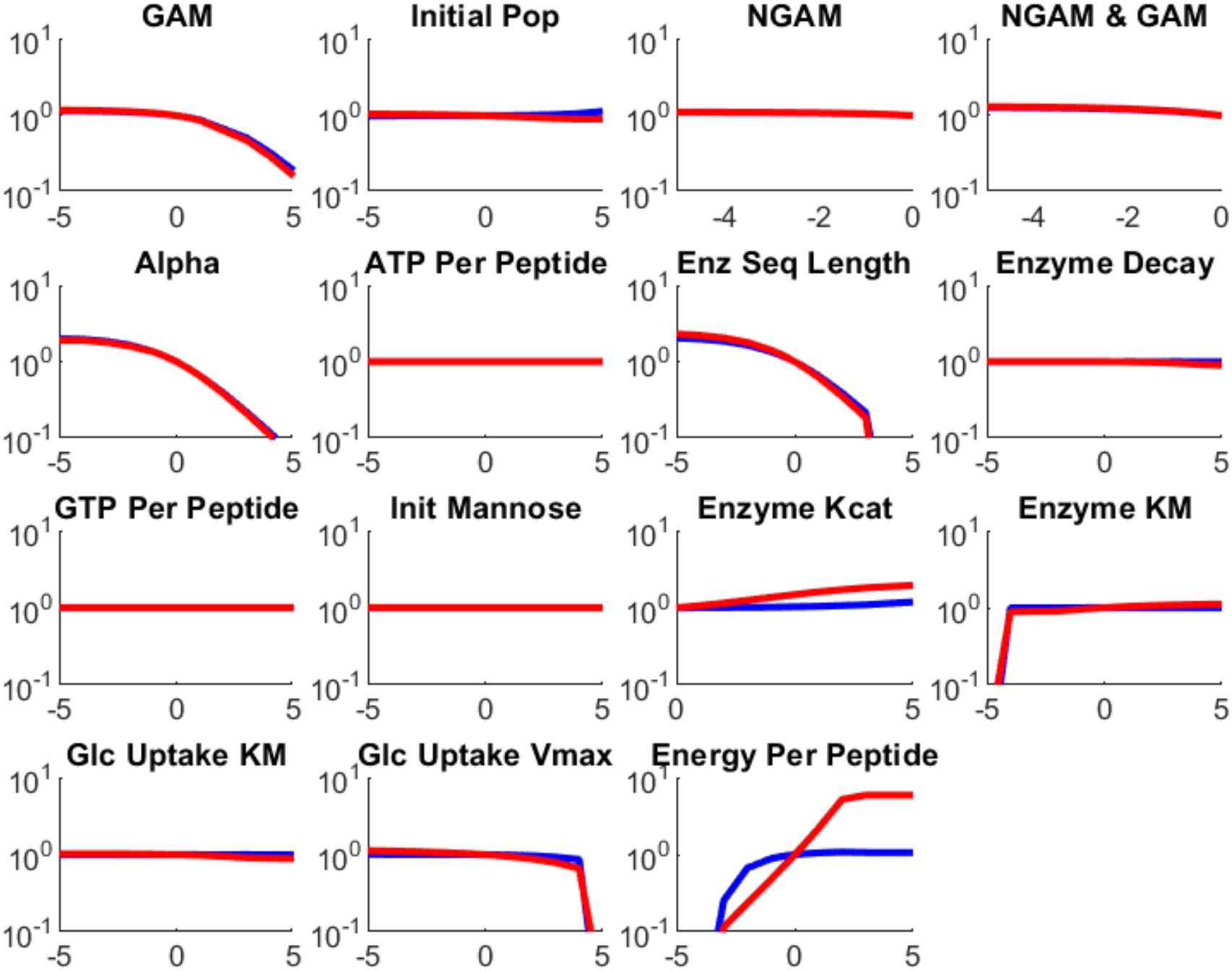
Sensitivity to Simulation Parameters.

Sensitivity of homogenous simulations to parameter values. Blue series indicate fold-change in biomass yield. Red series indicate fold-change in maximum growth rate.

Homogenous simulations were run as above, with each of the following parameters varied on a scale from 2^−5^ to 2^5^ times the original value: α, initial population, growth and non-growth associated maintenance reaction rate, ATP and/or GTP cost per peptide bond in the synthesis of the enzyme protein, enzymatic turnover rate, and Michaelis constant KM, maximum glucose uptake rate, half-saturation concentration for glucose uptake, enzyme decay rate, and the stoichiometric cost of production for a fixed amount of enzyme. Default values are listed in Tables S1, S2, and S3.

**Table S1:** Parameters used in the metabolic model.

| Parameter | Value | Units | Citation |
| --- | --- | --- | --- |
| $\alpha$ | 5.62e-4 | | |
| ATP per peptide (enzyme synthesis) | 8 |  | [88] |
| Cellulase $k_{cat}$ | 32 | $s^{-1}$ | [61] |
| Cellulase $K_M$ | 0.7 | $mmol/cm^3$ | [62] |
| Cell death rate | 0 | $h^{-1}$ | |
| Glucose per unit cellulose | 218.2 |  | [83] |
| Glucose uptake $K_M$ | 5.6e-4 | $mmol/cm^3$ | |
| Glucose uptake $V_{max}$ | 3.9 | $mmol/(gDW \cdot h)$ | |
| Growth Associated Maintenance (GAM) flux lower bound | 23.49 |  |  |
| GTP per peptide (enzyme synthesis) | 4 |  | [88] |
| Initial biomass | 1.35e-4 | gDW |  |
| Intracellular reaction flux lower bound | -1000 |  |  |
| Intracellular reaction flux upper bound | 1000 |  |  |
| Mannose uptake $K_M$ | 5.6e-4 | $mmol/cm^3$ | |
| Mannose uptake $V_{max}$ | 3.9 | $mmol/(gDW \cdot h)$ | |
| Non-Growth Associated Maintenance (NGAM) flux lower bound | 0.7 |  |  |
| Uptake reaction $K_M$ | 0.01 | $mmol/cm^3$ | |
| Uptake reaction $V_{max}$ | 10 | $mmol/(gDW \cdot h)$ | |

**Table S2:** Parameters for COMETS simulations.

| Parameter | Value | Units |
| --- | --- | --- |
| Allow Cell Overlap | TRUE |  |
| Allow Flux Without Growth | TRUE |  |
| Exchange Style | Monod |  |
| Extracellular Reaction Substeps | 10 |  |
| Objective Style | Max_Objective_Min_Total |  |
| Space Width | 4.6416 | cm |
| Time Step | 1 | hour |
| Enzyme Decay Rate | 1 | % per hour |

**Table S3:** Initial Media Composition.

| Metabolite | Millimoles |
| --- | --- |
| Ca<sup>2+</sup> | 10 |
| Cellulose | 0.01248 |
| Cl | 10 |
| CO<sub>2</sub> | 10000 |
| Cu<sup>2+</sup> | 10 |
| Ergosterol | 0.00252 |
| H<sub>2</sub>O | 10000 |
| H<sup>+</sup> | 1000 |
| K | 1000 |
| D-Mannose | 0.01598 |
| Mg<sup>2+</sup> | 10 |
| Mn<sup>2+</sup> | 10 |
| Na<sup>+</sup> | 10 |
| NH<sub>4</sub> | 10000 |
| O<sub>2</sub> | 0 |
| Oleate | 0.10557 |
| Palmitoleate | 0.10557 |
| PO<sub>4</sub><sup>3-</sup> | 10000 |
| SO<sub>4</sub><sup>2-</sup> | 10000 |
| Zn<sup>2+</sup> | 10 |

**Table S4:**
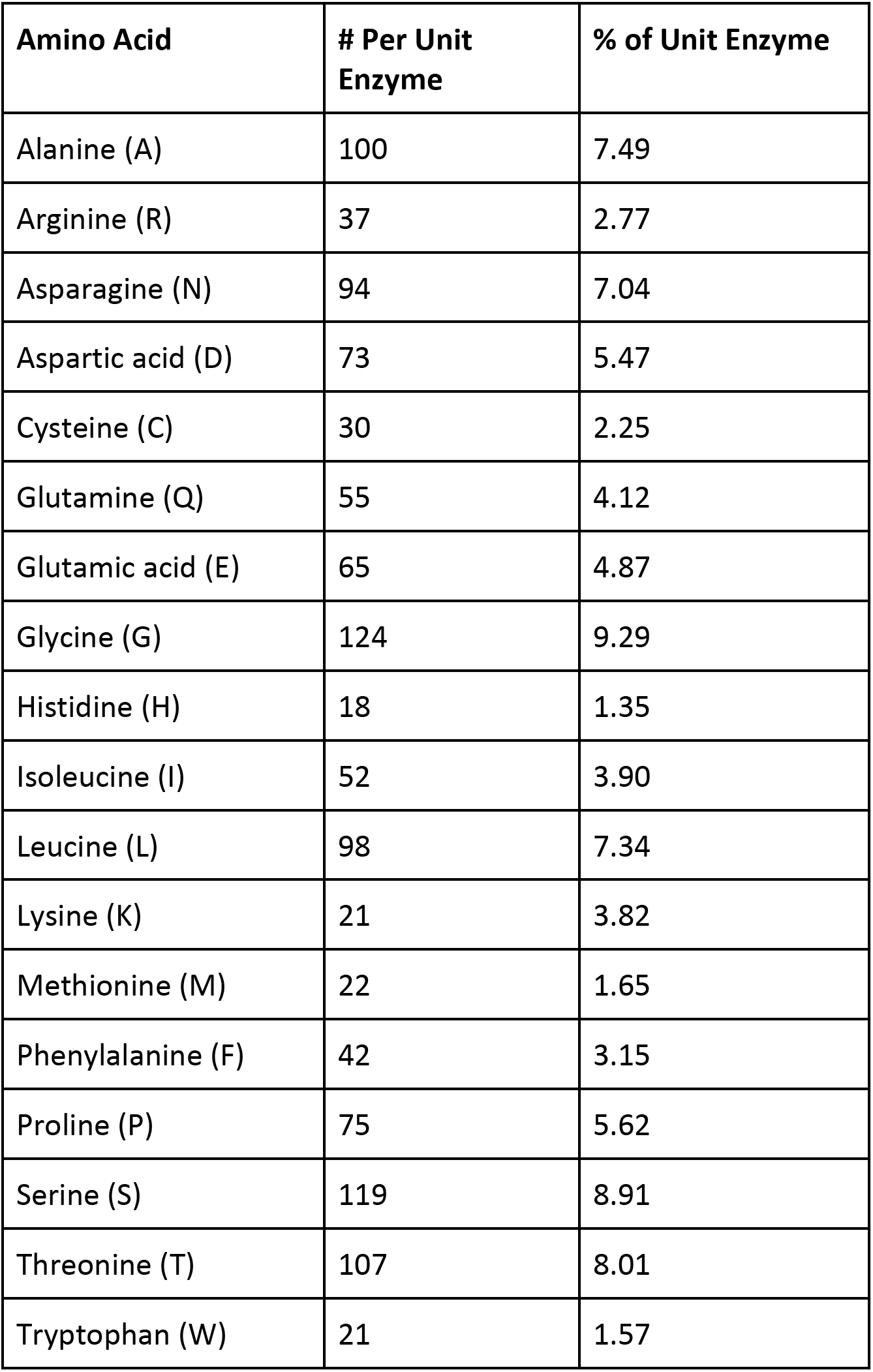
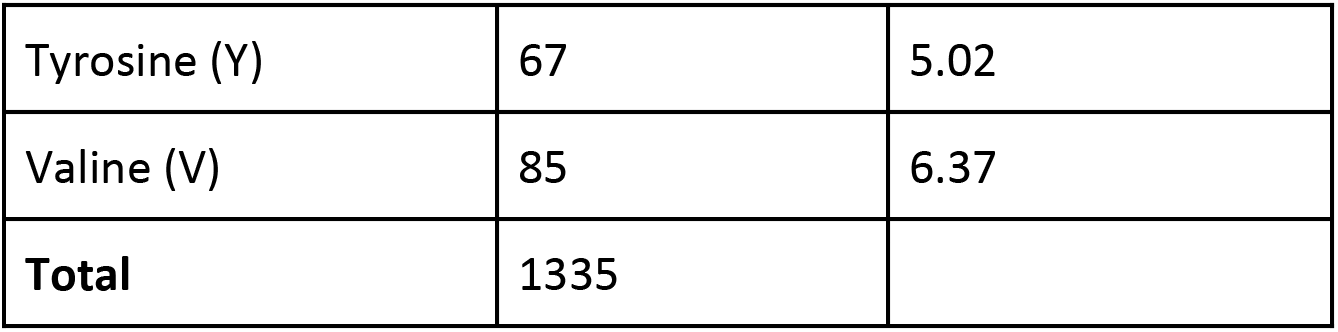
Enzyme Composition.

| Amino Acid | # Per Unit Enzyme | % of Unit Enzyme |
| --- | --- | --- |
| Alanine (A) | 100 | 7.49 |
| Arginine (R) | 37 | 2.77 |
| Asparagine (N) | 94 | 7.04 |
| Aspartic acid (D) | 73 | 5.47 |
| Cysteine (C) | 30 | 2.25 |
| Glutamine (Q) | 55 | 4.12 |
| Glutamic acid (E) | 65 | 4.87 |
| Glycine (G) | 124 | 9.29 |
| Histidine (H) | 18 | 1.35 |
| Isoleucine (I) | 52 | 3.90 |
| Leucine (L) | 98 | 7.34 |
| Lysine (K) | 21 | 3.82 |
| Methionine (M) | 22 | 1.65 |
| Phenylalanine (F) | 42 | 3.15 |
| Proline (P) | 75 | 5.62 |
| Serine (S) | 119 | 8.91 |
| Threonine (T) | 107 | 8.01 |
| Tryptophan (W) | 21 | 1.57 |
| Tyrosine (Y) | 67 | 5.02 |
| Valine (V) | 85 | 6.37 |
| **Total** | 1335 |  |

The stoichiometry of the amino acid components in the enzyme production reaction was set according to sequences obtained for the *Trichoderma reesei* endoglucanase Cel7b and the Saccharomycopsis fibuligera beta-glucosidase BGL1. This table displays the total amino acid proportions shown below.

